# Positive feedback and feedforward loops between PERIANTHIA, WUSCHEL-RELATED HOMEOBOX5 and GRF-INTERACTING FACTOR 1 modulate gene expression and function in the Arabidopsis root

**DOI:** 10.1101/439851

**Authors:** Adam Paul Fisher, Natalie Minako Clark, Rosangela Sozzani

## Abstract

The Arabidopsis root meristem consists of populations of stem cells that surround the mitotically less active cells known as the Quiescent Center (QC). The QC maintains the stem cells in a non-cell-autonomous manner through the function of the transcription factor (TF) WUSCHEL-RELATED HOMEOBOX5 (WOX5), which is required for columella stem cell (CSC) maintenance. However, whether WOX5 has a regulatory role in any other adjacent stem cells is less understood. To this end, we identified a set of TFs downstream of WOX5 in both QC and Cortex Endodermis Initial (CEI) cells. We then utilized Gene Regulatory Network (GRN) inference to identify GRF-INTERACTING FACTOR 1 (GIF1) as a key gene involved in positive feedback and feedforward loops with WOX5 as well as another stem cell regulator, PERIANTHIA (PAN). Finally, we constructed an ordinary differential equation model based on this inferred GRN to simulate GIF1, PAN, and WOX5 expression over time, which suggests the precise temporal expression of WOX5 and GIF1 is important to sustain QC function.

## Introduction

Stem cells divide to regenerate themselves and to generate all of the cell- and tissue-types in a multicellular organism. The continued ability to sustain stem cell maintenance within their micro-environment, the stem cell niche (SCN), is an important developmental characteristic that ensures proper tissue growth. In the *Arabidopsis thaliana* root SCN, different populations of stem cells are maintained by the Quiescent Center (QC), which generate short-range signals to repress differentiation (van den Berg et al., 1997). A known QC- derived signal is the homeobox transcription factor (TF) WUSCHEL-RELATED HOMEOBOX 5 (WOX5), which function to repress differentiation of one of the adjacent stem cell populations, namely the Columella Stem Cells (CSCs) (Petricka et al., 2012; Sarkar et al., 2007). Specifically, non- cell- autonomous WOX5 maintenance of CSCs takes place through the repression of the differentiation factor CYCLING DOF FACTOR 4 (Pi et al., 2015). However, whether WOX5 is expressed in, or moves to, the other stem cell populations, including the Cortex-Endodermis Initials (CEIs), is not known.

Recently, GRF-INTERACTING FACTOR 1 (GIF1) was shown to regulate the expression of SCARECROW (SCR) (Ercoli et al., 2018), which along with SHORT-ROOT, the D-type Cyclin CYCLIND6;1 (CYCD6:1) and RETINOBLASTOMA-RELATED (RBR) control the Cortex-Endodermis Initial (CEI) divisions to generate the cortical and endodermal tissue layers (Cui et al., 2007; Gallagher and Benfey, 2009; Gallagher et al., 2004; Helariutta et al., 2000; Levesque et al., 2006; Nakajima et al., 2001; Sozzani et al., 2010). In addition to regulating SCR, it was shown that GIF1 positively regulates WOX5 and that, together with additional GRF family members, GIF1 plays a role in QC function (Ercoli et al., 2018). However, whether GIF1 function is dependent on WOX5 regulation and whether GIF1 has a regulatory role outside the QC, is less understood.

In this study, we combine cell-type specific gene expression data with computational methods to better understand whether WOX5 and its downstream genes may play a role outside the QC and specifically in the CEI. To this end, we transcriptionally profiled QC cells in wild-type and *wox5-1* roots. We found that a number of differentially expressed (DE) genes in the QC are known to regulate CEI division, which led us to transcriptionally profile CEI cells in wild-type and *wox5-1* mutant backgrounds. Through differential gene expression analysis, GRN inference, and network motif analysis, we found that GIF1 acts in a positive feedback loop with WOX5 in both the QC and CEI. In support of a role for GIF1 in the QC and CEI, we showed that *gif1* mutants have disorganized QC cells and earlier periclinal CEI daughter (CEId) divisions. Next, we inferred a GRN using spatiotemporal gene expression data, which predicted that PAN regulates WOX5 and GIF1 in a positive feedforward loop. Finally, we built an Ordinary Differential Equation (ODE) model of this network that incorporates experimentally determined WOX5 and GIF1 oligomeric state. Our ODE model results suggest that PAN regulates WOX5 and GIF1 as a homodimer and that the *pan* mutant phenotype correlates with a temporal delay in WOX5 and GIF1 expression. Overall, our results suggest that positive feedback and feedforward loops between PAN, WOX5, and GIF1 modulate temporal expression of WOX5 and GIF1 to properly sustain QC activity.

## Results

### Analysis of WOX5 downstream genes in QC and CEI cells

To identify the genes regulated by WOX5 in the QC, we acquired gene expression profiles from the QC (cells expressing pWOX5:erGFP) of *wox5-1* and wild-type (WT) roots (see Methods, Figure 1A). We identified 2,407 differentially expressed (DE) genes (201 TFs) between *wox5-1* and WT backgrounds. Specifically, 935 genes (63 TFs) were significantly downregulated and 1,472 genes (138 TFs) were significantly upregulated in the *wox5* mutant, suggesting those genes are positively and negatively regulated by WOX5, respectively Supplemental Tables 1, 2). Within this list of 201 DE TFs, we found a number of genes known to regulate root development (Table 1), which included WIP DOMAIN PROTEIN 4 (WIP4), shown to be important for root initiation, as well as, ARABIDOPSIS HOMOLOG of TRITHORAX1 (ATX1/SDG27), shown to control root growth by regulating cell cycle duration. Notably, as a downstream gene of WOX5, we also found HEAT SHOCK TRANSCRIPTION FACTOR B4 (HSFB4)/ SCHIZORIZA (SCZ), which has been known to have a functional role in both QC as well as CEI cells. This finding led us to hypothesize that WOX5 may have a regulatory role in the CEI cells.

**Figure 1.**
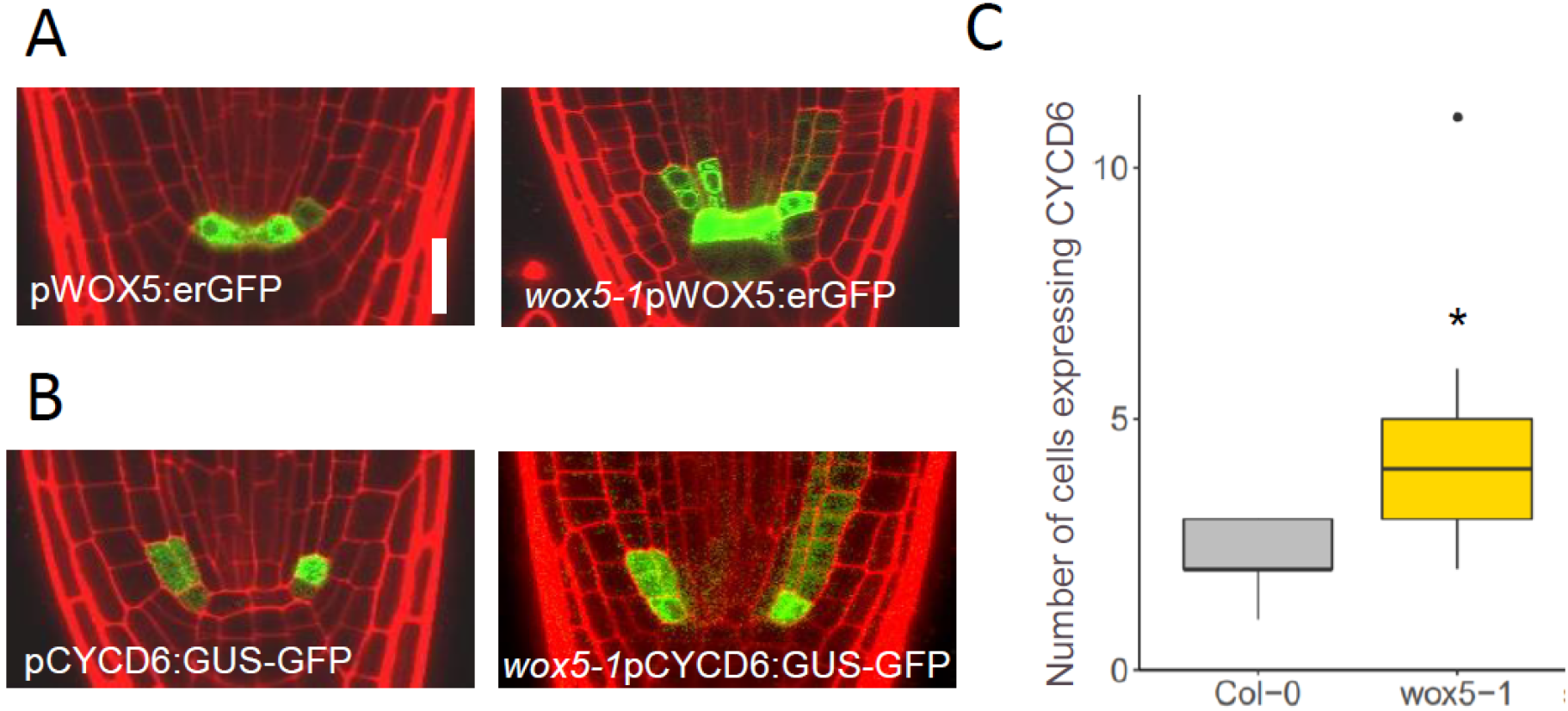
Expression of QC and CEI markers in the *wox5-1* mutant. (A) Representative images of roots expressing pWOX5:erGFP, which marks the QC cells, in Col-0 (left) and wox5-1 (right) backgrounds. Scale bar: 20 μm (B) Representative images of roots expressing pCYCD6:GUS-GFP, which marks the CEI, in Col-0 (left) and wox5-1 (right) backgrounds. (C) Quantification of the number of cells expressing pCYCD6:GUS-GFP in WT and *wox5-1* mutants (n=19 and n=15, respectively). * denotes p < 0.05, Wilcoxon test. The solid dot represents outliers.

**Table 1.** List of a 7 transcription factors (TFs) previously shown to regulate some process in root development

| Gene Name | References | Function |
| --- | --- | --- |
| PLT4/BBM | Galinha et al., 2007 | Promote root formation |
| WIP4 | Crawford et al., 2015 | Initiation of the root meristem when combined in *wip4wip5ntt* triple mutant |
| ATX1/SDG27 | Napsucialy-Mendivil et al., 2014 | Root growth and cell production |
| HSFB4/SCZ | ten Hove et al., 2010 | Controls separation of stem cell fate |
| AN3/GIF1 | Ercoli et al., 2018 | Control root meristem homeostasis when combined in *an3gif2* or *an3gif2gif3* |
| HAM2 | Engstrom et al., 2011 | Root growth and cell divisions |

To test if WOX5 may regulate genes in the CEI, we first examined the expression of the CEI marker CYCLIND6;1:GFP-GUS (hereafter referred to as CYCD6) (Sozzani et al., 2010) in *wox5-1* roots. We found that significantly more cells express CYCD6 compared to WT, suggesting WOX5, through a yet unknown mechanism, is required to limit CYCD6 expression to the CEI and supports a functional role for WOX5 in CEI division (p<0.05, Wilcoxon test, Figure 1C). To identify genes downstream of WOX5 in the CEI, we performed cell-type specific gene expression profiling of the CEI (cells expressing CYCD6) in *wox5-1* and WT roots (Figure 1B). We found 2,302 DE genes/133 TFs, of which 1,295 genes/65 TFs were significantly downregulated, and the remaining 1,007 genes/68 TFs were significantly upregulated in the *wox5-1* mutant. This result further suggests that WOX5 has a regulatory role in the CEI (Supplemental Tables 3, 4). Importantly, we found that 16 out of 133 TFs DE in the CEI were also DE in the QC dataset. Taken together, the changes in gene expression between WT and *wox5-1* roots, along with the expansion of the CEI-specific marker in a *wox5-1* mutant, suggest that WOX5 plays a role in both the QC and CEI.

### WOX5 regulates QC function and CEI daughter division through GIF1

To determine if WOX5 regulates genes similarly in the QC and CEI, we focused on the 200 genes (16 TFs) identifies as DE in both the QC and CEI of *wox5-1* roots compared to WT (Supplemental Table 5, 6). Moreover, to better understand the relationship between the 16 DE TFs, we inferred a Gene Regulatory Network (GRN) using RTP-STAR as inference algorithm (de Luis Balaguer, 2018; Shibata et al., 2018). The application of RTP-STAR with our QC and CEI cell-type resulted in an inferred network of 12 TFs (Figure 2A), for which we performed a network motif analysis to rank the functionally important factors. We specifically focused on network motifs that were previously demonstrated to contain factors that are important for development (Alon, 2007; Ingram et al., 2006; Milo et al., 2002). We added the number of times a gene appears in each of these motifs and normalized the sum to calculate the Normalized Motif Score (NMS) for each gene in the network (see Methods). Given the well-characterized developmental role of WOX5 in the root, we used the NMS of WOX5 as the cutoff for significance and selected the 7 genes with higher importance scores than WOX5 for further characterization (Table 2). Among these 7 TFs were HSFB4/SCZ, DECREASE WAX BIOSYNTHESIS (DEWAX), ZAT18, HISTONE DEACETYLASE 2C (HD2C), MYB DOMAIN PROTEIN 34 (MYB34), an unnamed bHLH family TF (At2g22760), and ANGUSTIFOLIA3 (AN3)/GRF-INTERACTING FACTOR1 (GIF1). Interestingly, these TFs showed a similar expression pattern, either activated or repressed in both the QC and CEI between *wox5-1* and WT roots, suggesting a similar regulatory role of WOX5 in these two different cell types.

**Figure 2.**
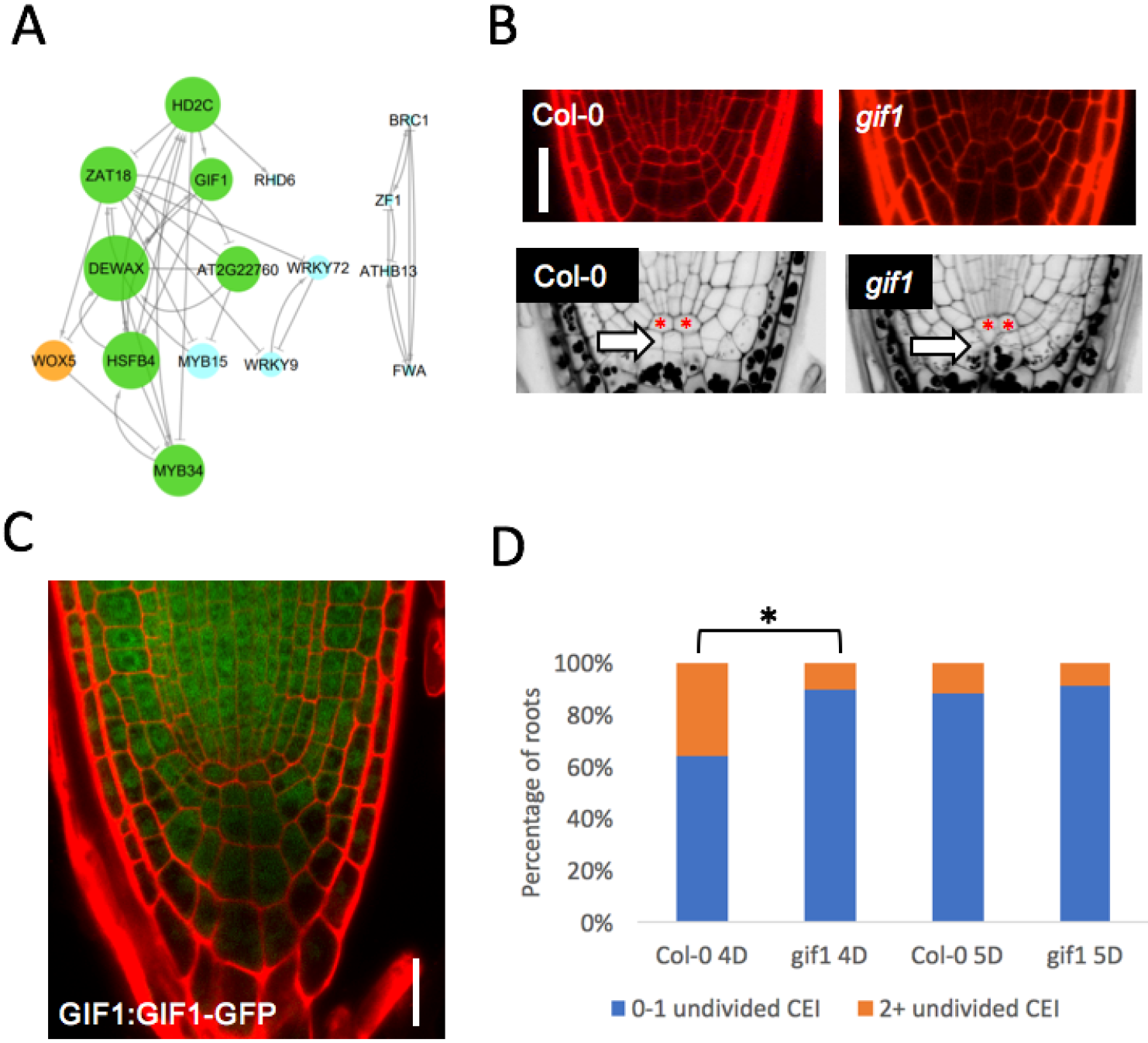
GRN inference identifies GIF1 as a regulator of QC and CEI maintenance. (A) Inferred gene regulatory network of genes DE in both QC and CEI. In the WOX5 subnetwork, the size of the node corresponds to normalized motif score (NMS) Green nodes were selected for downstream analysis. Arrows represent positive regulation, horizontal lines negative regulation, and circles unknown signs. (B) (Top row) Representative images of wild-type and *gif1* mutants. Scale bar = 20 μm.(Bottom row) Representative images of mPSPI stained roots of wild-type and *gif1* mutant. The red asterisks mark the QC, and the white arrow denotes the first layer of columella cells. (C) GIF1:GIF1-GFP translational fusion expression in the root meristem. (D) Quantification of the percent of roots with low (0-1 cells, blue) and high (2+ cells, orange) numbers of undivided CEI in Col-0 and *gifl* plants. * denotes p < 0.05, Fisher’s Exact Test.

**Table 2.**
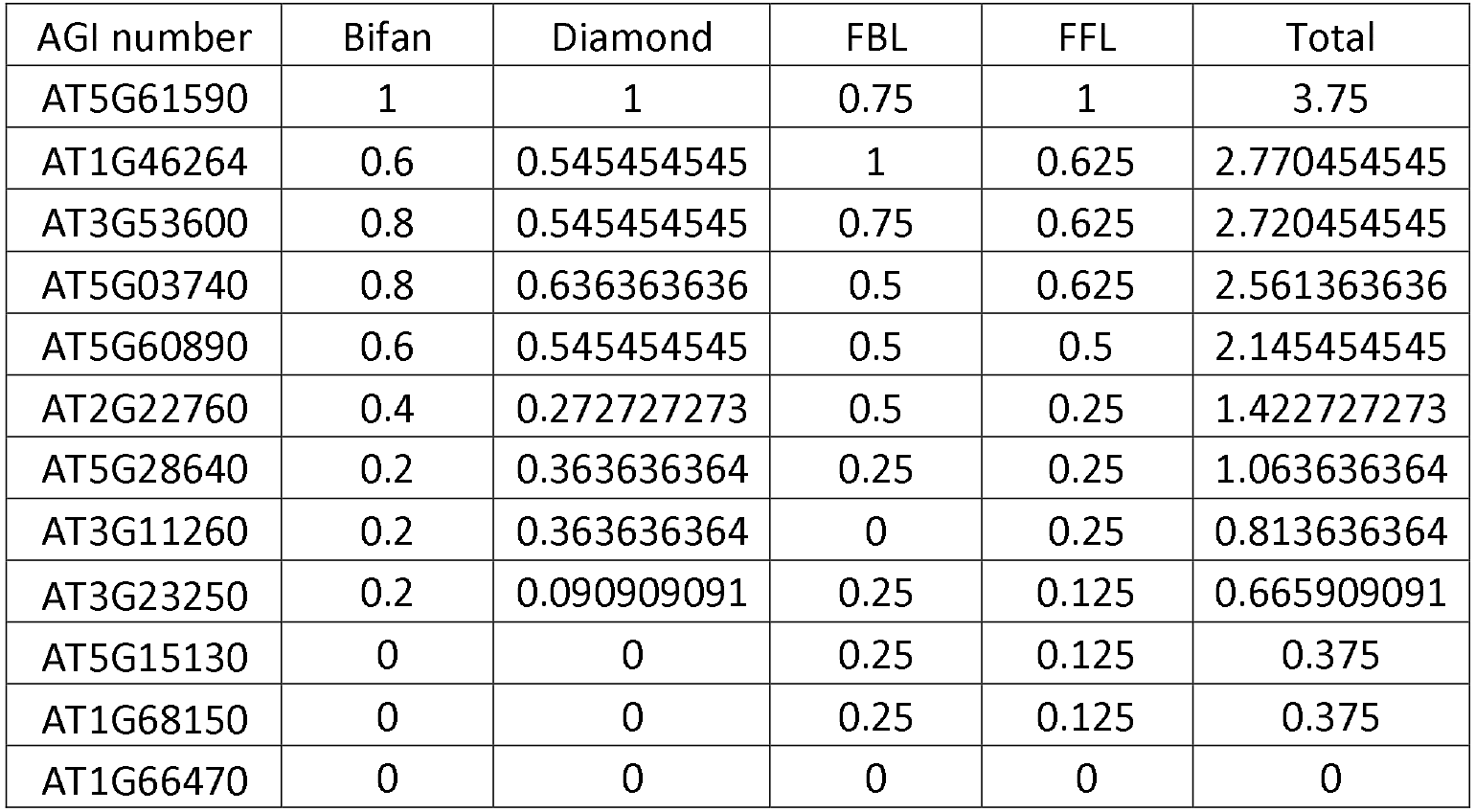
List of the 12 transcription factors inferred by RTP-STAR and normalized motif score for the 4 different network motifs. Italicized font indicates those transcription factors that were further investigated.

| AGI number | Bifan | Diamond | FBL | FFL | Total |
| --- | --- | --- | --- | --- | --- |
| AT5G61590 | 1 | 1 | 0.75 | 1 | 3.75 |
| AT1G46264 | 0.6 | 0.545454545 | 1 | 0.625 | 2.770454545 |
| AT3G53600 | 0.8 | 0.545454545 | 0.75 | 0.625 | 2.720454545 |
| AT5G03740 | 0.8 | 0.636363636 | 0.5 | 0.625 | 2.561363636 |
| AT5G60890 | 0.6 | 0.545454545 | 0.5 | 0.5 | 2.145454545 |
| AT2G22760 | 0.4 | 0.272727273 | 0.5 | 0.25 | 1.422727273 |
| AT5G28640 | 0.2 | 0.363636364 | 0.25 | 0.25 | 1.063636364 |
| AT3G11260 | 0.2 | 0.363636364 | 0 | 0.25 | 0.813636364 |
| AT3G23250 | 0.2 | 0.090909091 | 0.25 | 0.125 | 0.665909091 |
| AT5G15130 | 0 | 0 | 0.25 | 0.125 | 0.375 |
| AT1G68150 | 0 | 0 | 0.25 | 0.125 | 0.375 |
| AT1G66470 | 0 | 0 | 0 | 0 | 0 |

To test if any of these TFs have a role in the QC and/or CEI function, we examined mutants of each of these genes for developmental phenotypes, such as a disorganized stem cell niche and/or aberrant ground tissue patterning. Accordingly, *hsfb4/scz* mutant was previously shown to misexpress multiple QC-specific markers and lack the undifferentiated CSC (ten Hove et al., 2010). Of the remaining 6 TFs, no obvious morphological phenotypes were observed in *dewax, zat18, hd2c, myb34,* and *at2g22760* roots 5 days post stratification (DPS) (Supplemental Figure 1). However, approximately 56% (25/45 roots) of *gif1* mutant roots showed a disorganized stem cell niche (Figure 2B). In addition, we found that *gif1* mutants contained starch granules in the cells that are normally CSC, suggesting that GIF1 plays a role in CSC maintenance (Figure 2B). To determine if GIF1 plays also a role in CEId divisions, we compared the number of undivided CEId cells present in *gif1* and WT roots at 4 and 5 DPS. We classified the roots into two groups, those with low numbers (0-1) and those with high numbers (2+), of undivided CEId. We found that WT roots had a significantly higher number of undivided CEId cells at 4 DPS compared to *gif1* mutants (35.71% in WT vs 10.34% in *gif1* mutants, p<0.05, Fisher’s Exact test), suggesting that more CEI divisions occur in the *gif1* mutant. However, we did not see a significant difference in the number of undivided CEI cells at 5 DPS (11.90% in WT vs 8.57% in *gif1* mutants, Fisher’s Exact Test). Taken together, these results suggest that GIF1 regulates the timing of CEI division and supports a role for GIF1 in stem cell maintenance downstream of WOX5.

### PAN and WOX5 regulate GIF1 through positive feedback and feedforward loops

We previously observed that mutations in PERIANTHIA (PAN) showed a similar phenotype to *wox5* roots, so we sought to understand if this phenotype could be due to the regulation of similar factors such as GIF1. PAN was previously identified as QC-enriched gene, and when we examined the gene expression profile of QC cells in *pan* mutant plants, we found that GIF1 is among the 4465 DE downstream genes of PAN (de Luis Balaguer et al., 2017) (Supplemental Table 7). Further, when we compared the transcriptomic profiles of QC cells collected from *wox5-1* and *pan* roots, we found that, in addition to GIF1, WOX5 was among the 589 common DE genes (28 shared DE TFs) (Supplemental Table 8). This led us to hypothesize that in the QC, PAN could regulate WOX5, and both WOX5 and PAN could regulate GIF1. To predict how PAN and WOX5 regulate their common downstream targets in a spatiotemporal manner, we built a gene regulatory network of PAN that includes WOX5 and the 27 additional TFs shown to be differentially expressed in both the *pan* and *wox5-1* roots. We first used RTP-STAR (de Luis Balaguer, 2018; Shibata et al., 2018) on our QC-specific, steady-state, gene expression data to build a network that contained the predicted QC-specific regulations at 5 DPS. Next, we inferred a second network using GENIST, a pipeline developed to use time course data to infer dynamic networks (de Luis Balaguer et al., 2017). To incorporate the regulation between PAN, WOX5, and their downstream targets on a time scale that would reflect regulations taking place in a developmental context, we used a time course gene expression dataset obtained from root meristems collected every 24 hours from 3 to 7 DPS (de Luis Balaguer et al., 2017). Lastly, to predict a GRN that incorporates both QC-specific and temporal gene regulation, we combined the two networks obtained from these pipelines.

The resulting GRN combining both inferred networks from RTP-STAR and GENIST contains 26 of the 28 shared DE TFs (Figure 3A). Importantly, this network recapitulates the three previously described regulations observed in the *wox5-1* and *pan* mutants, specifically PAN regulation of WOX5, PAN regulation of GIF1, and WOX5 regulation of GIF1 (Figure 3A, Supplemental Tables 1,7). Moreover, we used the NMS to quantify gene importance, and we found that PAN, GIF1, and WOX5 are all ranked in the top 50% of genes (Supplemental Table 9). To determine the sign of regulation between PAN, WOX5, and GIF1, we examined how each gene changed in either mutant gene expression dataset. We found that all three of the regulations are positive, thus, forming a positive feedforward loop. In addition, our network predicts that GIF1 also activates WOX5, forming a positive feedback loop. To validate this network prediction, we examined the pWOX5:erGFP in a *gif1* mutant background and found that WOX5 expression was significantly lower compared to WT (Figure 3B, C). Taken together, the inferred GRN using spatiotemporal gene expression data led us to hypothesize that the regulation between PAN, WOX5, and GIF1 could have a role in sustaining QC function, specifically for CSC and CEI maintenance.

**Figure 3.**
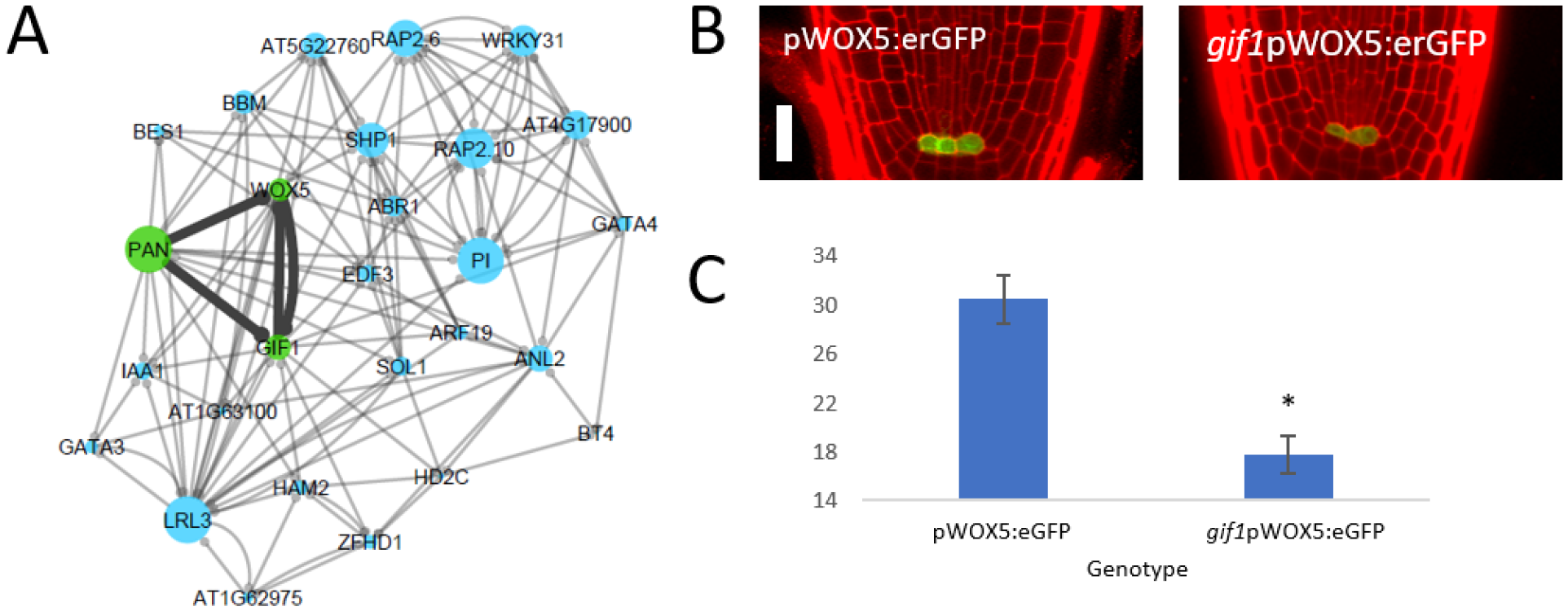
Gene regulatory network inference predicts a PAN-WOX5-GIF1 regulatory module in the QC. (A) Gene regulatory network of genes differentially expressed in both *wox5-1* and *pan* mutants inferred by RTP-STAR and GENIST. The regulation between PAN, WOX5, and GIF1 (green) is highlighted (black edges). Circles represent an unknown sign of regulation. Size of the nodes represents NMS score. (B) Representative confocal images roots expressing pWOX5:eGFP in WT (left) and *gifl* mutant background (right). Scale bar = 20 μm (C) Corrected total cell fluorescence (CTCF) of *gifl* pWOX5:eGFP (n=22) and pWOX5:eGFP (n=22). * denotes p < 0.05, two-tailed f-test.

### Mathematical modeling predicts that PAN regulation is important for proper timing of WOX5 and GIF1 expression

To determine the dynamics of the PAN-WOX5-GIF1 regulatory network, we built a mathematical model of Ordinary Differential Equations (ODEs). This ODE model includes the regulations predicted by our network, namely, the positive regulation of WOX5 and GIF1 by PAN, the positive regulation of WOX5 by GIF1, and the positive regulation of GIF1 by WOX5. Given that our network did not predict any upstream regulation of PAN, we assumed PAN expression was constant over time. This resulted in generating ODEs only for GIF1 and WOX5. To determine which parameters of the model were most important, we performed a global sensitivity analysis (see Methods). The sensitivity analysis identified the production and degradation terms, as well as the oligomeric states of GIF1 and WOX5, as the most important parameters of the model whose values needed to be estimated or experimentally determined (Supplemental Tables 10, 11, Supplemental Figure 2). Accordingly, we experimentally determined the oligomeric states of GIF1 and WOX5 using scanning Fluorescence Correlation Spectroscopy (scanning FCS). Specifically, we performed Number and Brightness (N&B) on Arabidopsis roots expressing either pGIF1:GIF1:GFP or pWOX5:WOX5:GFP translational fusion. We found that both GIF1 and WOX5 primarily exist as a monomer (98.67% and 96.01%, respectively) with a very small amount of dimer (1.33% and 3.99%, respectively) (Supplemental Figure 3). Thus, we fixed the oligomeric state of GIF1 and WOX5 as a monomer in our ODE model.

The oligomeric state of PAN was also identified as an important parameter in our model, and we used our model to estimate this parameter. To do this, we changed the parameter for PAN oligomeric state and then fit the model prediction to the *wox5* and *pan* mutant transcriptomic data, as we reasoned that the best parameter value will be able to recapitulate the transcriptomic changes seen in these mutants. We first simulated WT plants by setting the sensitive parameter values so that GIF1 and WOX5 reach the expression levels measured in 5 DPS WT plants after time (t) = 24 arbitrary units (AU) (Supplemental Table 12). We then assumed that PAN existed as a monomer and tested whether we could recapitulate the *pan* mutant data by adjusting the value of PAN expression as observed in the mutant. We found that reducing PAN in the model incorporating only PAN monomer did not cause the reduction of WOX5 and GIF1 that was experimentally observed in the *pan* mutant (Supplemental Figure 4).

This led us to hypothesize that PAN may need to be in a higher oligomeric state to recapitulate our experimental data (e.g gene expression in roots of mutants). Thus, we adjusted our model so PAN could form a dimer (Figure 4A, Methods). Using this modified model, we found we could accurately reduce the levels of WOX5 and GIF1 in a *pan* mutant simulation when all (100%) of PAN exists as a dimer. In addition, this model accurately recapitulated the reduction of GIF1 in a *wox5-1* mutant background (Figure 4A). Therefore, our model predicted that PAN needs to exist as a dimer to regulate WOX5 and GIF1.

**Figure 4.**
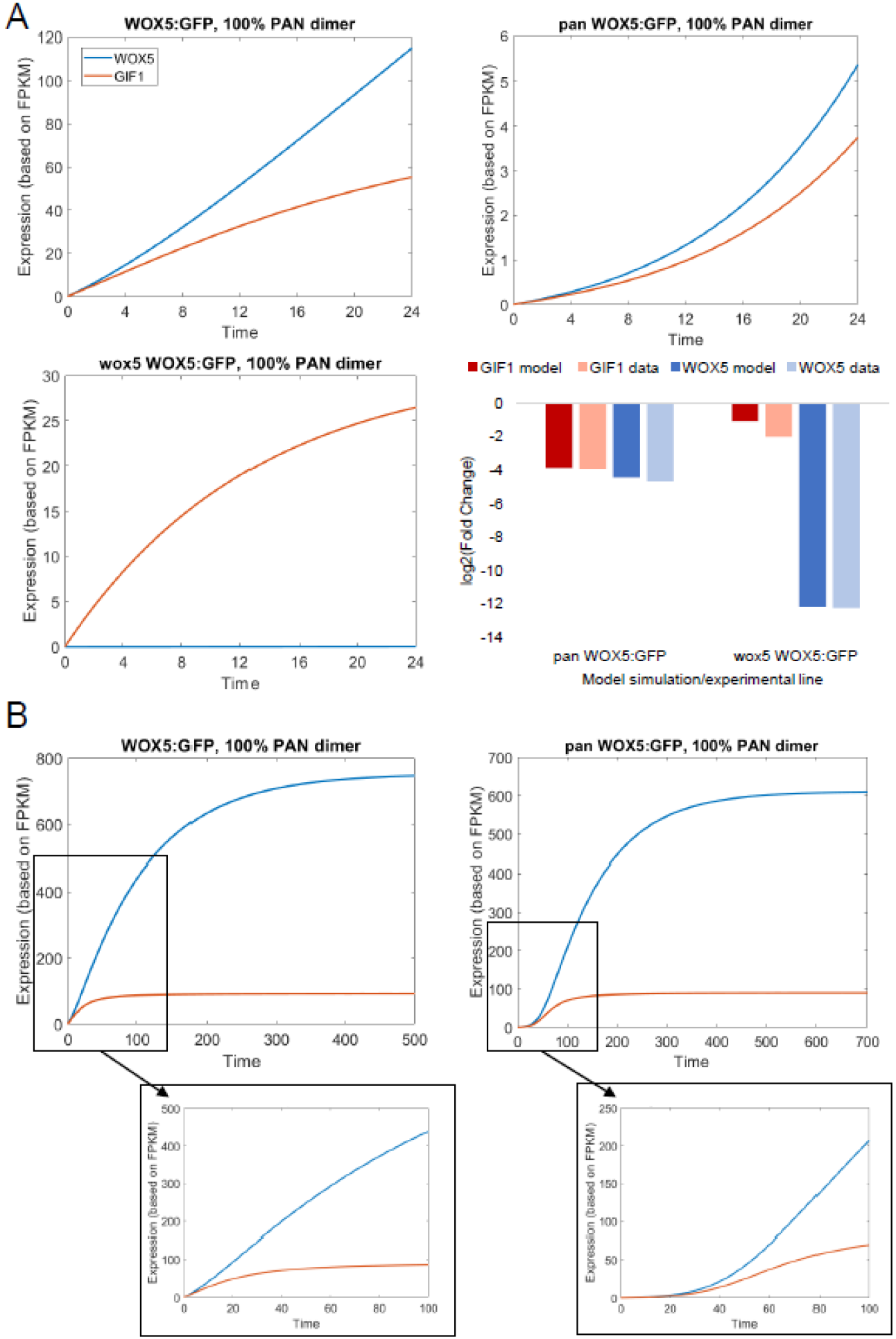
Mathematical modeling quantifies WOX5 and GIF1 expression over time. (A) Model simulation of WOX5:GFP (top left), *pan*WOX5:GFP (top right), and *wox5-1*WOX5:GFP (bottom left) for 24 time points. The blue line is WOX5 expression, the red line is GIF1 expression. (Bottom right) Quantification of fold change predicted by the model (dark bars) compared with data (light bars) at t=24. (B) Model simulation of WOX5:erGFP (left) and panWOX5:erGFP (right) showing when the expression levels reach steady state. Inset: t=0 to t=100.

Finally, we used our model to predict how changes in PAN and WOX5 expression affect the dynamics of gene expression. We found that in the WT simulation, GIF1 reached its steady state value around t = 80, while WOX5 reaches its steady state at a much later time (around t = 500) (Figure 4). In a *wox5* mutant simulation, GIF1 still reached its steady state around t = 80 (Supplemental Figure 5), suggesting that WOX5 does not affect the timing of GIF1 expression but rather the overall levels. Thus, our model prediction suggested that in *wox5-1* plants a reduction in steady-state levels of GIF1 expression is caused by the elimination of both the positive feedback and feedforward loops (Figure 5). In contrast, in a *pan* mutant simulation, both genes took significantly longer to reach their steady states. GIF1 did not reach its steady state until t = 200 (150% increase), while WOX5 finally reached steady state at t = 700 (40% increase). Additionally, the WOX5 steady state in the *pan* mutant was slightly lower (29.1% change) than in wild type (Figure 4A), suggesting that WOX5 and GIF1 expression eventually reach their WT levels but at a much slower rate. Therefore, our model predicted that in the *pan* mutant, the loss of the positive feedforward loop causes a delay in WOX5 and GIF1 expression (Figure 5). Taken together, our model predicts that the overall levels of GIF1 and WOX5, as well as how quickly they reach steady-state expression levels, which in turn would regulate QC maintenance.

**Figure 5.**
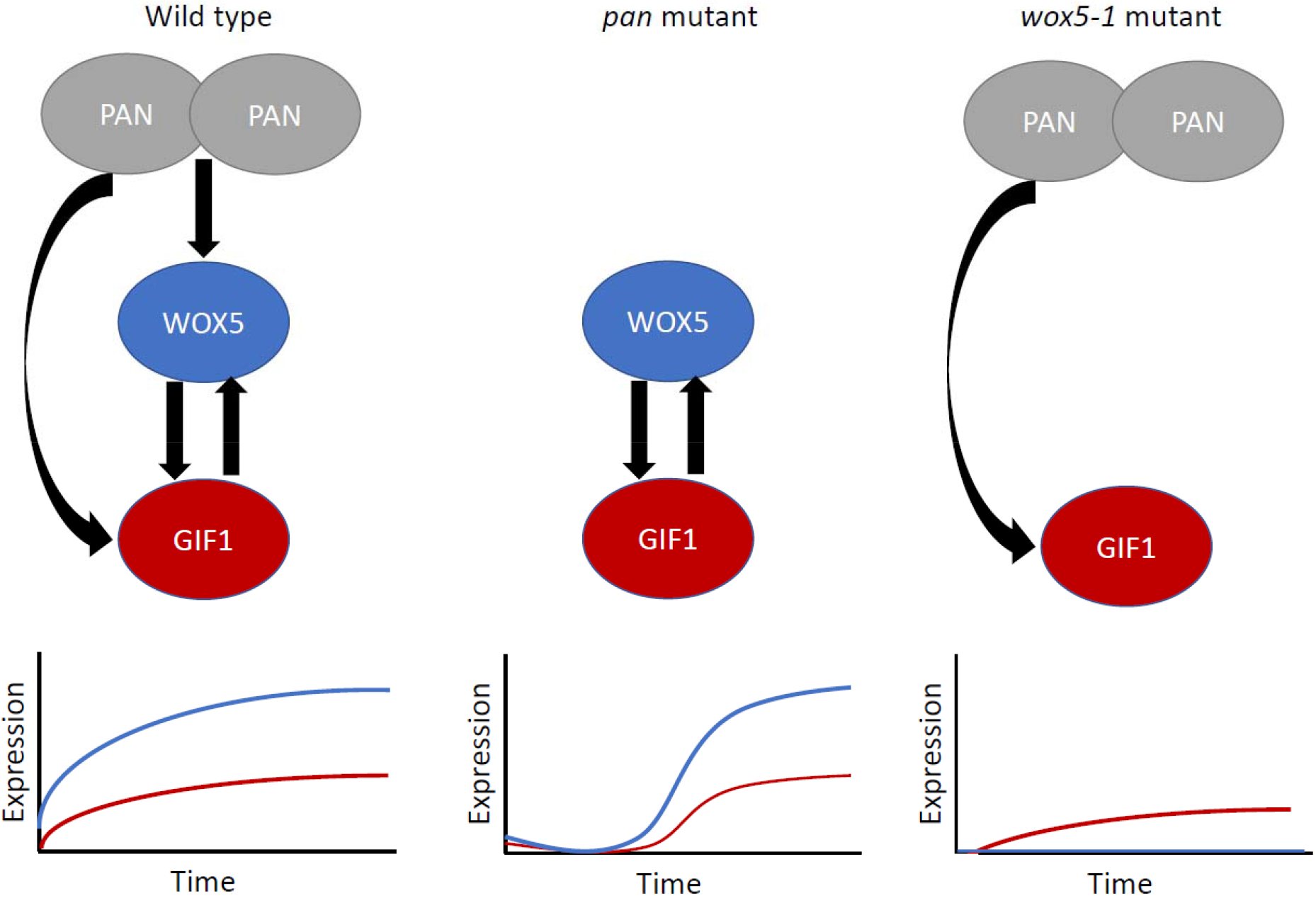
Model simulation of the PAN-WOX5-GIF1 regulatory network. In wild-type plants (left), PAN homodimerizes to regulate both WOX5 and GIF1, which both reach a steady state over time. In a *pan* mutant (center), WOX5 and GIF1 still reach their steady states, but with a significant time delay. In a *wox5-1* mutant (right), the expression of both WOX5 and GIF1 is significantly reduced.

## Discussion

Our results, which include cell-type specific gene expression, gene regulatory network inference, and ODE modeling, allowed us to better understand the regulation between PAN, WOX5, and GIF1. Our model included numerous parameters such as production rates, degradation rates, and oligomeric states. A global sensitivity analysis revealed that the oligomeric states are important for the model prediction. Other models have similarly shown that oligomeric states are important parameters, such as a model of SHR and SCR in the endodermis (Clark et al, 2016). Further, this sensitivity analysis allowed us to predict that PAN must exist as a homodimer to recapitulate the mutant expression phenotypes. The resulting model suggests that in absence of PAN expression, WOX5 and GIF1 increase at a slower rate than in the WT simulation but do still eventually reach their WT steady-state levels (Figure 5). The change in WOX5 and GIF1 expression is the result of the loss of the positive feedforward loop, showing how this regulatory motif modifies gene expression over time. Further, this result could possibly explain why the observed *pan* phenotype is 12% penetrant (de Luis Balaguer et al., 2017), as stochastic differences in gene expression between mutant roots could determine if the increase in WOX5 and GIF1 is “slow enough” to cause a developmental phenotype. The model also suggests that in the absence of WOX5 expression, GIF1 levels never reach their WT steady-state levels due to the loss of both the positive feedback and feedforward loops (Figure 5). This could help explain the *wox5-1* root phenotype as both *gif1* and *wox5-1* mutants have overall stem cell niche disorganization. However, while a *wox5-1* mutant has increased expansion of the CEI marker CYCD6, the patterning of cortex and endodermis is otherwise normal. Because we found that CEI cells in the *gif1* mutant undergo premature division, this result suggests a role of GIF1 in regulating CEI divisions. Despite the fact that GIF1 regulates CEI divisions through a yet unknown mechanism, it is tempting to speculate that GIF1 could be involved in a gene network containing feedback and feedforward similarly to what was described in this study.

## Methods

### Plant Material and Growth Conditions

The pWOX5:erGFP (Sarkar et al., 2007), pWOX5:WOX5-GFP (Pi et al., 2015), *wox5-1* (Sarkar et al., 2007), pCYCD6:GUS-GFP (Sozzani et al., 2010), *myb34 (Celenza et al., 2005), dewax* (Sparks et al., 2016), and *gif1* (Kim and Kende, 2004) were previously described. The remainder of the lines were ordered from the Arabidopsis Biological Resource Center (ABRC), *zat18* (Salk_077654, CS358533), *hd2c* (CS846589, Salk_129799), and *at2g22760* (CS444213) (Alonso et al., 2003). *wox5-1*pWOX5:erGFP and *wox5-1*pCYCD6:GUS-GFP lines were generated by crossing the lines described above.

Seeds used for confocal microscopy were surface sterilized using fumes produced by the combination of a solution containing 50% bleach and 1M hydrochloric acid (HCl). The seeds were subsequently stratified and imbibed in sterile water for 2 days at 4C. After 2 days, the seeds were plated on 1X Murashige and Skoog (MS) medium supplemented with 1% sucrose and grown vertically in long day conditions (16 hours light / 8 hours dark) at 22C. All plant roots were imaged at 5 days after stratification. For cell-type specific transcriptomic profiling experiments, seeds were wet sterilized using 50% bleach, then 100% ethanol, followed by 7 water rinses. Seeds were stratified and imbibed in sterile water for 2 days at 4C. After 2 days, the seeds were plated on Nitex mesh squares on top of 1X MS medium supplemented with 1% sucrose and grown vertically at 22C in long-day conditions as described above.

### Cell-type specific transcriptomic profiling on marker lines

Three hundred to five hundred mg of pWOX5:erGFP, *wox5-1* pWOX5:erGFP, pCYCD6:GUS-GFP, and *wox5-1*pWOX5:erGFP seeds were wet sterilized and plated (see Plant Materials and Growth Conditions) for each biological replicate. After 5 days of growth, approximately 1mm of the root tip was collected and protoplasted as described (Birnbaum et al., 2005). GFP positive cells were collected using a MoFlo cell sorter into a vial containing a solution of beta-mercaptoethanol and RLT buffer. RNA was extracted using the Qiagen RNeasy Micro kit. Libraries were prepared using the SMART-Seq v3 Ultra Low RNA Input Kit for Sequencing and the Low Library Prep Kit v1 from Clontech. Libraries were sequenced on an Illumina HiSeq 2500 with 100 bp single-end reads.

### RNA-seq data analysis

Reads from all experiments, including the previously published *pan* pWOX5:eGFP data (de Luis Balaguer et al, 2017) were filtered using ea-utils, mapped using RSubread (Liao et al., 2013), and FPKM values acquired using Cufflinks (Trapnell et al., 2012). Differential expression analysis was performed using PoissonSeq (Hu et al., 2012) with a cutoff of q < 0.05 and fold change > 2 in the pWOX5:eGFP vs *wox5-1*pWOX5:eGFP and pWOX5:eGFP vs *pan*pWOX5:eGFP comparisons, while a cutoff of q < 0.2 and fold change > 2 was used in the pCYCD6:GUS-GFP vs *wox5-1*pCYCD6:GUS-GFP comparison. The latter q-value was changed to reflect the fact that those cell samples were transcriptionally similar and to achieve a similar number of differentially expressed genes in both the CEI and QC. (GEO number: GSE118102)

### Confocal Microscopy

Confocal microscopy was completed using a Zeiss LSM 710. A 488nm laser was used for green channel acquisition, and a 570nm laser was used for red channel acquisition. A 10uM propidium iodide solution was used to stain cell walls for visualization. mPS-PI staining to visualize starch granules was performed as described in (de Luis Balaguer et al., 2017). For Number and Brightness (N&B) acquisition, 12-bit raster scans of a 256 × 256 pixel region of interest were acquired with a 100nm pixel size and 12.61μs pixel dwell time as described in (Clark and Sozzani, 2017). Heptane glue was used during the N&B acquisition to prevent movement of the root sample as described in (Clark and Sozzani, 2017).

### Normalized Motif Score

Four different motifs were used to calculate the normalized motif score, namely feedforward loops, feedback loops, diamond, and bi-fan motifs (Alon, 2007; Ingram et al., 2006; Milo et al., 2002). All motifs were significantly enriched in our network compared to a randomly generated network of the same size. We counted the times each gene appeared in each motif, normalized the counts to a scale of 0 to 1, and added the normalized counts together to calculate the Normalized Motif Score (NMS) for each gene. The most functionally important genes are those that have high NMS scores.

### Number and Brightness (N&B) Analysis

N&B analysis was performed using the SimFCS software (Clark and Sozzani, 2017; Digman et al., 2005). The 35S:GFP line was used to normalize the autofluorescence/background region of the image (S-factor of 2.65) and determine monomer brightness (brightness of 0.26). A 128×128 region of interest was used on all images to measure oligomeric state specifically in the QC.

### Mathematical model formulation

Two Ordinary Differential Equations (ODEs) measuring the change in WOX5 (W) and GIF1 (G) expression over time were built for the mathematical model. The first equation, measuring the
change in WOX5 expression over time, contains a Hill equation that incorporates PAN (P) activation of WOX5 and GIF1 activation of WOX5. The second equation, measuring the change in GIF1 expression over time, contains a similar Hill equation that incorporates PAN and WOX5 activation of GIF1. The oligomeric states of all 3 proteins are unknown parameters, and both WOX5 and GIF1 are assumed to degrade linearly. Given there is no predicted upstream regulation of PAN in our network, PAN is treated as a constant value.

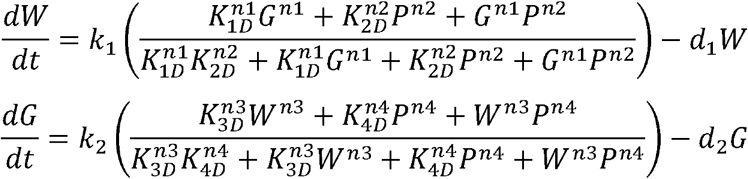

After determining that the PAN oligomeric state is an important parameter (see Sensitivity analysis, Supplemental Figure 2, Supplemental Tables 10,11), we modified the equations such that a certain proportion of PAN (P_2_) could exist as a dimer. These modified equations also incorporate the experimental results that WOX5 and GIF1 exist as monomers (Supplemental Figure 3).

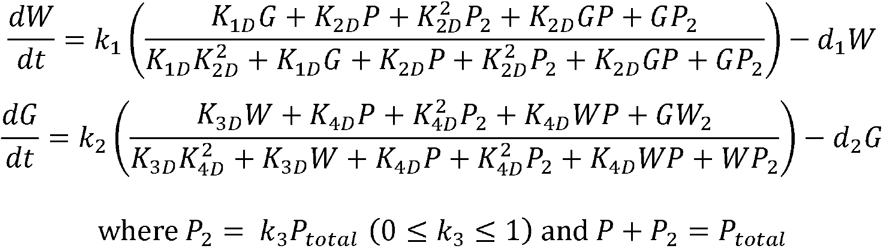

### Sensitivity analysis of mathematical model and parameter estimation

The total Sobol index (Clark et al., 2016; Saltelli et al., 2010; Sobol, 2001) was used to calculate the sensitivity of the model parameters. For each of 10 technical replicates, 1000 values were sampled for each model parameter from a uniform distribution. The model was evaluated for each parameter set, and then the total Sobol index was calculated for each technical replicate. The 10 technical replicates were then averaged and used for statistical analysis. The oligomeric states for GIF1 (*n*_*1*_) and WOX5 (*n*_*3*_) were experimentally determined (Supplemental Figure 3), and the remaining sensitive parameters were estimated using gene expression data.

## Author Contributions

Conceptualization, A.P.F., N.M.C and R.S.; Data Acquisition and Analysis, A.P.F., and N.M.C, Mathematical Model N.M.C, Writing, Review & Editing, A.P.F., N.M.C and R.S.

## Acknowledgements

We thank Dr. Kensuke Kawade for *gif1* and pGIF1:GIF1:GFP seeds, Dr. Thomas Laux for pWOX:WOX5-GFP seeds, and Dr. John Celenza for MYB34 seeds. We also thank the Arabidopsis Biological Resource Center (ABRC) for proving seeds. APF is supported by a NSF GRFP (DGE-1252376). NMC is supported by a NSF GRFP (DGE-1252376). RS is supported by a NSF CAREER grant (MCB-1453130). The authors have declared that no competing interests exist.

---

**Supplemental Figure 1:**
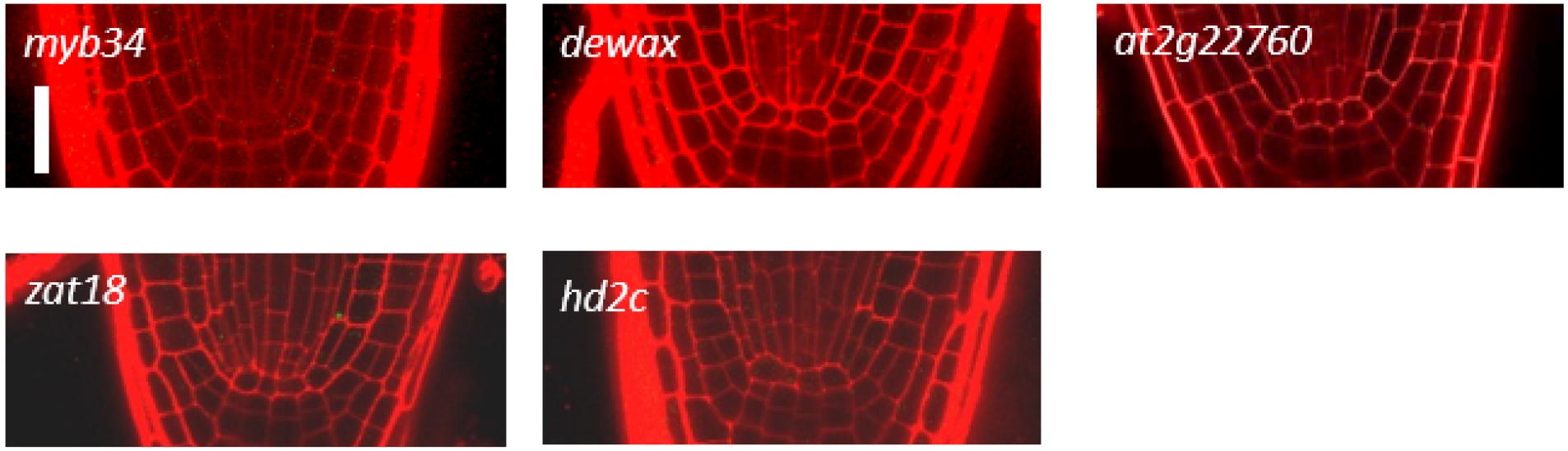
Stem cell niche morphology of mutants of 5 TFs examined

**Supplemental Figure 2:**
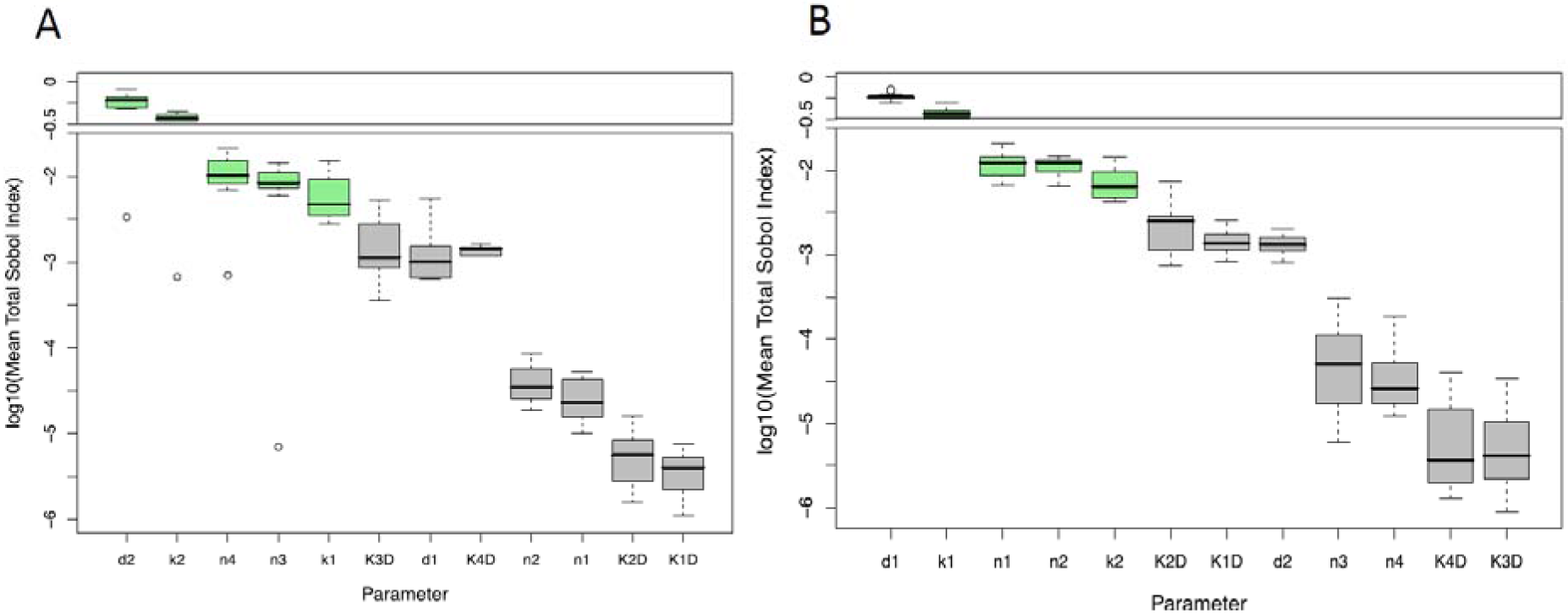
Sensitivity analysis of the mathematical model for (A) the GIF1 equation and (B) the WOX5 equation. Green boxes represent the sensitive parameters (p<0.05, Wilcoxon test with Steel Dwass for multiple comparisons)

**Supplemental Figure 3:**
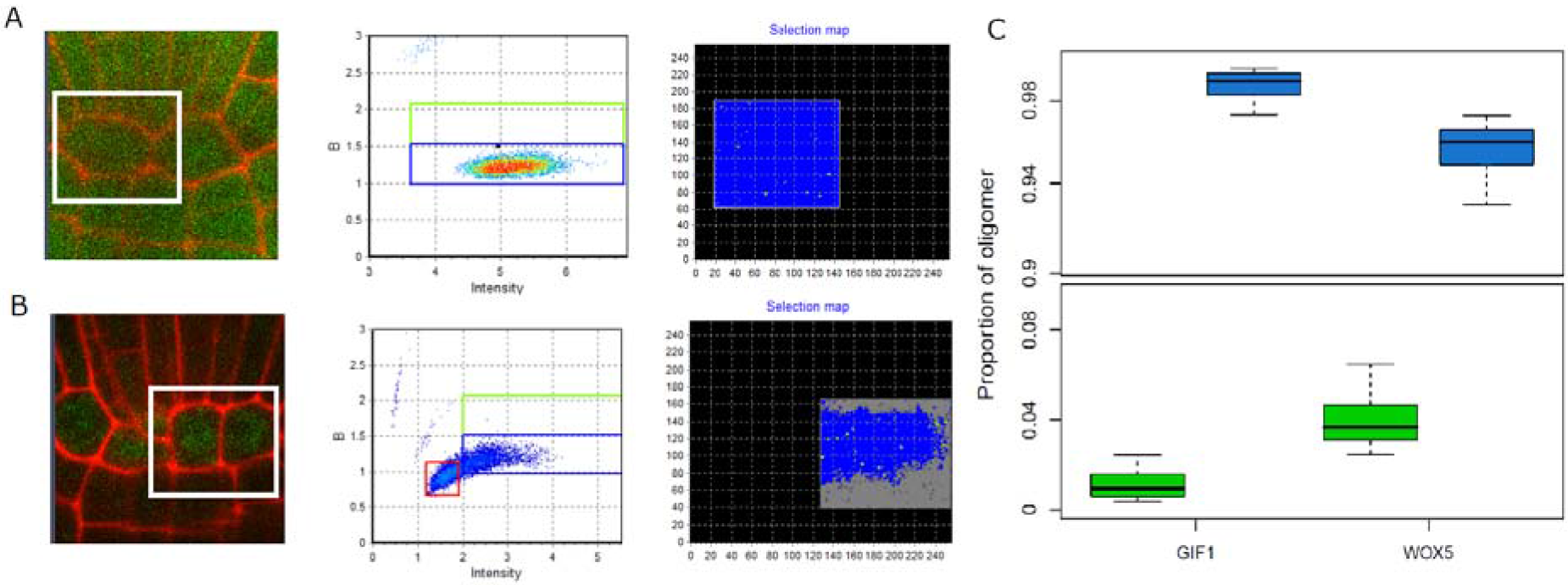
N&B analysis on GIF1 (A) and WOX5 (B) translational fusions. (A,B) show a representative image (left), brightness vs intensity plot (middle), and false-colored map (right). The white box represents a 128×128 pixel region of interest used for analysis. In the false-colored image, grey represents background, blue represents monomer, and green represents

**Supplemental Figure 5:**
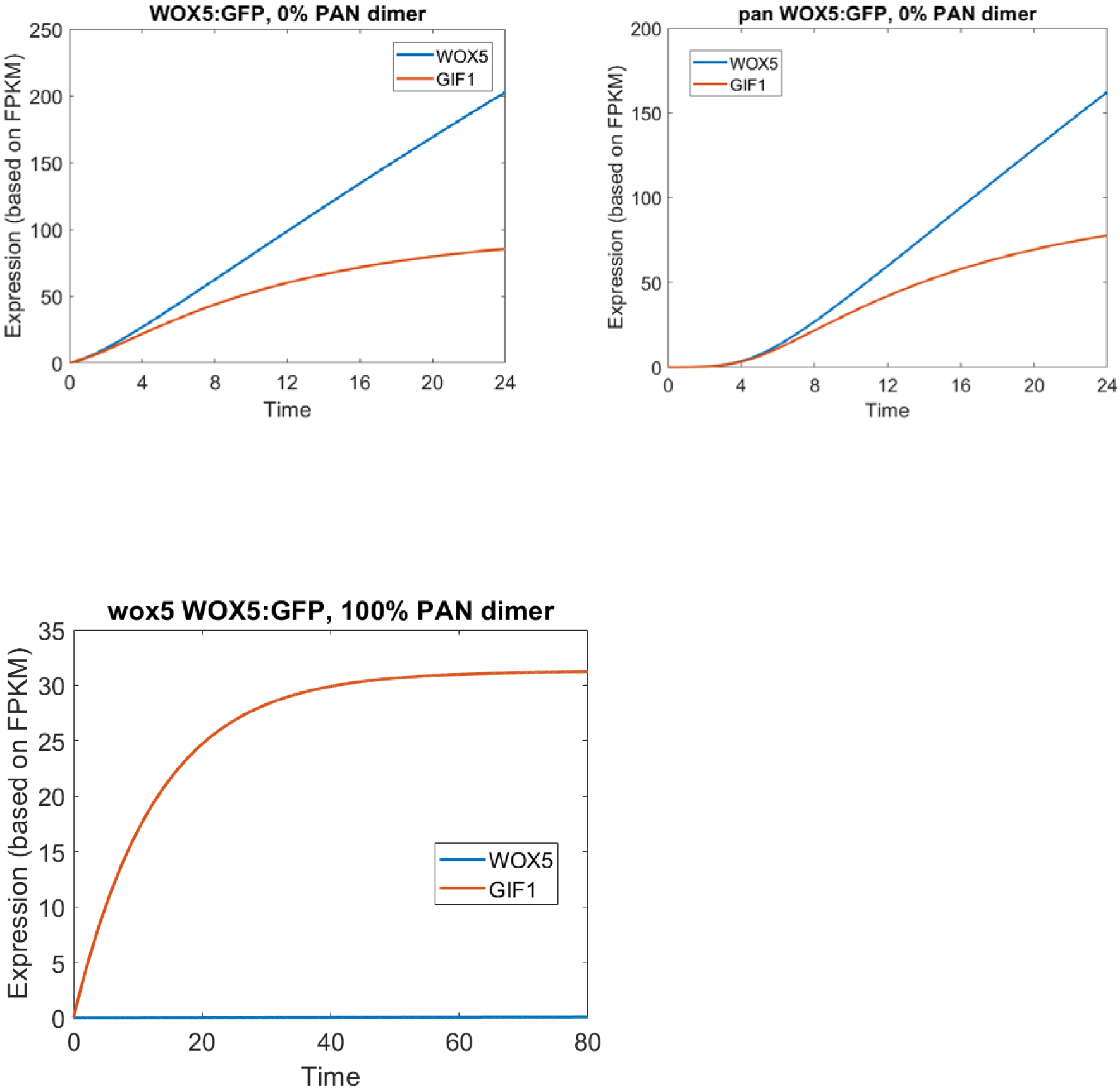
Model simulation of *wox5-1* mutant showing when GIF1 expression reaches steady state at t=80.

**Supplemental Table 1:**List of 2,407 differentially expressed genes between cells expressing pWOX5:eGFP in *wox5-1* and wild-type backgrounds.

**Supplemental Table 2:**List of 201 differentially expressed transcription factors between cells expressing pWOX5:eGFP in *wox5-1* and wild-type backgrounds.

**Supplemental Table 3:**List of 2,302 differentially expressed genes between cells expressing pCYCD6::GUS-GFP in *wox5-1* and wild-type backgrounds.

**Supplemental Table 4:**List of 133 differentially expressed transcription factors between cells expressing pCYCD6::GUS-GFP in *wox5-1* and wild-type backgrounds.

**Supplemental Table 5:**List of 200 differentially expressed genes in common between cells expressing pWOX5:eGFP and pCYCD6::GUS-GFP in *wox5-1* mutant and wild-type background.

**Supplemental Table 6:**List of 16 differentially expressed transcription factors in common between cells expressing pWOX5:eGFP and pCYCD6::GUS-GFP in *wox5-1* and wild-type background.

**Supplemental Table 7:**List of 4,465 differentially expressed genes between cells expression pPAN:GFP in *pan057190* and wild-type backgrounds.

**Supplemental Table 8:**List of 589 differentially expressed genes in common between cells expression pWOX5:eGFP in *wox5-1* and *pan057190* mutant backgrounds

**Supplemental Table 9:**Normalized motif score of 28 shared TFs between cells expression pWOX5:eGFP in *wox5-1* and *pan057190* mutant backgrounds

**Supplemental Table 10:**Sensitivity analysis for GIF1

**Supplemental Table 11:**Sensitivity analysis for WOX5

**Supplemental Table 12:**Model parameter values

